# Pharmacological inhibition of titan-like cells formation suggests that accumulation of endogenous free radicals that correlate with mitochondrial changes are required to induce this transition in *Cryptococcus neoformans*

**DOI:** 10.1101/2022.04.06.487428

**Authors:** Irene García-Barbazán, Rocío García-Rodas, Martin Sachse, Daniel Luque, Diego Megías, Oscar Zaragoza

**Affiliations:** Mycology Reference Laboratory. National Centre for Microbiology. Instituto de Salud Carlos III. Carretera Majadahonda-Pozuelo, Km 2. Majadahonda 28220. Madrid. Spain.; Electron Microscopy Unit. Central Core Facilities, Instituto de Salud Carlos III; Advanced Optical Microscopy Unit. Central Core Facilities, Instituto de Salud Carlos III

**Keywords:** *Cryptococcus neoformans*, off-patent drug repurposing, titan-like cells, reactive oxygen species, mitochondria

## Abstract

*Cryptococcus neoformans* is an encapsulated yeast able to cause disease (mainly meningoencephalitis) among immunosuppressed patients, mostly HIV+. This yeast can form the so-called titan cells *in vivo*, which are cells of an abnormal larger size due to an increase in both the capsule and the cell body size (total size reaching between 50-70 microns). This phenomenon can be partially reproduced *in vitro* to obtain cells of an intermediate size (25-30 um), which have been denominated titan-like cells. In this work, we have screened 1,520 compounds from the Prestwick Chemical Library and identified off-patent drugs that inhibited titan-like cell formation *in vitro*. We developed an automated fluorescence-based microscopy assay and identified 64 compounds as possible inhibitors of titan-like cells *in vitro*. We chose 10 of these compounds to confirm their inhibitory effect and confirmed them as inhibitors of titan-like cells with dose-response curves. Several of the compounds identified had antioxidant properties (i.e., retinoic acid), indicating a possible role of free radicals during titan cell formation. Using fluorescent probes, we found that there was an endogenous accumulation of ROS during cell growth, which was inhibited in the presence of retinoic acid. Furthermore, we found that during titanization, there were significant changes in the mitochondria, which is the main organelle where ROS are produced. We hypothesize that an intracellular increase of free radicals at the mitochondria might be a triggering signal to induce titanization.

**Importance:** *Cryptococcus neoformans* is an excellent model to investigate fungal pathogenesis. This yeast can produce “titan cells”, which are cells of an abnormal larger size that contribute to the persistence of the yeast in the host. In this work, we have used a new approach to characterize them, which is identifying drugs that inhibit this process. We have used a repurposing off- patent drug library, combined with an automatic method to image and analyse fungal cell size. In this way, we have identified many compounds that inhibit this transition. Interestingly, several compounds were antioxidants, allowing us to confirm that endogenous ROS and mitochondrial changes are important for titan cell formation. This work provides new evidences of the mechanisms required for titanization. Furthermore, the future characterization of the inhibitory mechanisms of the identified compounds by the scientific community will contribute to better understand the role of titan cells in virulence.

## Introduction

*Cryptococcus neoformans* is a pathogenic basidiomycete yeast widely distributed in the environment that can cause disease in humans (1). In the last years, its incidence has decreased in developed countries due to the antiretroviral therapy (ART), but it is still responsible for hundreds of thousands of deaths each year in developing areas (2, 3). Cryptococcosis affects mainly to immunocompromised patients, especially those with AIDS/HIV, being, after tuberculosis, the second cause of death in these patients (2, 3). Infection is initiated by inhalation of infective particles, which can colonize the lungs (4–6). Immunocompetent hosts control the infection, and yeasts are eliminated or remain as latent forms. In contrast, in people with weakened immune system, the yeasts can disseminate to the central nervous system, causing meningoencephalitis, which can be fatal without treatment (7).

*Cryptococcus neoformans* is a unique fungal pathogen due to its ability to adapt to different environments and infect many different hosts, including mammals (including some aquatic, as dolphins), plants, insects and amoebas (8–17). There are several reasons that explain this unique adaptation among fungal pathogens, such its ability to melanize (18, 19) or to survive within phagocytic cells (20–23). But one of the most important aspects is the presence of a polysaccharide capsule around the cell body that interferes with the host immune response (24–33). In addition, the capsule can increase in size in response to multiple factors *in vitro* and during infection (34–37), conferring protection to stress factors (38, 39).

Another striking characteristic of this pathogen during infection is its ability to dramatically increase its cell size (including both the capsule and the cell body, delimited by the cell wall), producing what has been denominated as titan cells (TC) which can reach up to 50-70 microns *in vivo* (40–43). Titan cells have been described as those cells with a cell body above 15 microns or with a total cell size (including capsule) above 30 microns (44). These cells are uninuclear polyploid cells with a big vacuole, a thick cell wall and a dense capsule, and its formation *in vivo* depends on signalling pathways and receptors such as PKA and Gpr4 respectively (42, 43). Due to their size, they can persist for longer periods in the host as they cannot be phagocytosed, and their presence can inhibit the phagocytosis of cells of regular size (45). The *in vivo* factors that trigger titan cell formation are poorly characterized, but a high proportion of titan cells have been found in mice with a Th2 polarized response or during asymptomatic infections (42, 46). In addition, coinfection of mice with MATa/MATα strains results in a significant higher proportion of titan cells in the lungs (43).

The process of titanization can be partially reproduced *in vitro* by incubating at different conditions, including low nutrient media supplemented with serum and a CO_2_ enriched atmosphere with oxygen limitation (47–49). This results in the appearance of cells of an intermediate size (around 15-25 microns) which had been denominated as titan-like cells. Titan-like cells are also induced by bacterial components, such as the peptidoglycan subunit muramyl dipeptide (47). The possibility to mimic the titan cells *in vitro* has allowed unravelling several signalling pathways and genes required for titan cell formation, such as PKA, PCK, and also the negative regulators, such as Tsp2, Usv101 (47–49).However, the exact molecular mechanisms involved in this process remain unknown.

In this work, we present a different strategy to identify new pathways and processes involved in titan-like cell formation. In particular, we have performed a screening to identify off-patent compounds that pharmacology inhibit titan-like cell development with the purpose of characterizing their effect on this process and to identify possible new targets, which could provide new insights in this process. Drug repurposing is a strategy that in the last years has gain great relevance to find new biological activities with clinical use. In the case of clinical mycology, drug repurposing has allowed to identify a large number of drugs with antifungal properties (50–60). We have used the Prestwick Chemical Library, which contains 1,520 off-patent drugs and we have identified a large number of drugs (around 60) that blocked titanization in *C. neoformans*. We noticed that several inhibitory drugs had antioxidant properties, so we hypothesized that an endogenous accumulation of free radicals in the cell might be one of the signals that triggers titan cell formation in *C. neoformans*.

## Materials and Methods

### Prestwick Chemical Library and selected compounds

The Prestwick Chemical Library (Prestwick Chemical Libraries, Domain Therapeutics, Strasbourg, Illkirch, France) was used to screen compounds that inhibit the formation of titan-like cells in *C. neoformans.* The library contains 1,520 off-patent compounds approved by different agencies as the Food and Drug Administration (FDA) and the European Medicines Agency (EMA). The Prestwick Chemical Library is prepared in 96-well plates, each well containing a different compound at 10 mM in 100% Dimethyl Sulfoxide (DMSO). Columns 1 and 12 in each plate were empty without compounds.

### Yeast strains and growth conditions

*Cryptococcus neoformans* H99 strain (var. *grubii*, serotype A, (61)) was used in all experiments. Yeasts were routinely grown in Sabouraud liquid or solid medium (Oxoid) at 30 °C. For the induction of titan-like cells, we followed the protocol described by (49). Briefly, the yeasts were grown overnight in Sabouraud liquid medium at 30 °C with moderate shaking at 150 r.p.m. The cells were then washed twice with 50 mM MOPS (Sigma Aldrich) and suspended at 2 x 10^4^ cells/mL in Titan Cell Medium (TCM) composed by 5% Sabouraud, 5% inactivated Fetal Calf Serum (FCS, Biological Industries) and 15 µM sodium azide (Sigma Aldrich), diluted in 50 mM MOPS adjusted to pH 7.3. Cells were incubated for 16 h at 37 °C with a 5% CO_2_ enriched atmosphere. In some experiments, titan-like cells were induced as described above in the presence of different DMSO (Sigma Aldrich) concentrations (ranging from 2% to 0.03%, two-fold serial dilutions). As control, cells in TCM without DMSO were carried out in parallel.

### Screening protocol and automated fluorescence-based microscopy protocol

To evaluate the effect of the compounds of the chemical library on titan-like cells formation, we first performed a 1/10 dilution of the compounds to obtain intermediate plates at 1 mM and 10% DMSO, in 96-well round bottom plates (Falcon). Five microliters from these intermediate plates were diluted 20 times in sterile distilled water to obtain 2x stocks of the compounds (0.05 mM in 0.5% DMSO in a final volume of 100 µl) in 96-well plates with flat glass bottom (Greiner). In parallel, cells incubated o.n. in Sabouraud liquid medium as described above were prepared at 2 x 10^4^ cells/mL in 2x TCM, and 100 µl were added to each well containing the compounds. In this way, the final screening was performed in TCM with a cell density of 10^4^ cells/mL with the compounds at 25 µM and 0.25% DMSO. The following controls were added to columns 1 and 12 (four wells of each control in each plate): 1) cells grown in 1x TCM, 2) cells grown in 1x TCM without serum, 3) cells grown in sterile distilled water, and 4) cells grown in Sabouraud liquid medium. All the controls contained DMSO 0.25%.

The plates were incubated for 16 h at 37 °C with a 5% CO_2_ enriched atmosphere. After this incubation, all the wells were visually observed with a DMI3000 microscope (Leica microsystems) to identify inhibitory compounds of titanization. In addition, we also developed an automated protocol that allowed us to take pictures of all the wells and measure the cell size in the presence of all the compounds. We used lactofuchsin, a dye that binds to the surface of fungi. This dye provides a light red staining of the cells, which is strongly fluorescent and easily observed in fluorescence microscopes using the standards rhodamine filters. A lactofuchsin stock was prepared with acid fuchsin (Sigma) at 1 mg/mL in 63% of lactic acid (Merck), and from this stock we made a 1/30 dilution in distilled water (33 µg/mL in 2.1% of lactic acid). Finally, 30 µL from this dilution were added to each of the wells of the screening, so the final lactofuchsin concentration in each well was 4.3 µg/mL with lactic acid at 0.27%.

After staining, the cells were directly observed with a Cytell automatic microscope (GE Healthcare Life Sciences) using the 10x objective. Five pictures from each well were taken in bright field and with the rhodamine fluorescence filter. In total, 480 images were obtained for each 96-well plates. The images were analyzed in two different automatic ways. First, we used the analysis options from the Cytell software, so a dot plot representing the size of the cells detected and the fluorescence intensity was obtained for each well. Then, the 480 images were exported in TIF format (16-bits), and analyzed using FIJI software (62) using the Batch Mode option, with an in house designed macro. Briefly, this macro creates first a mask of the fluorescent cells and then, the feret diameter and area of each cell identified in the mask is determined. In this way, around 250-500 cells per well are automatically measured. Results were exported as a .csv document, which was further processed with Microsoft Excel program using the Pivot tables option, obtaining the area, average diameter and standard deviation of the cells in the presence of each compound.

### Dose response curves of selected compounds for confirmation

Selected compounds were bought as powder (Prestwick Chemical Libraries, Domain Therapeutics, Strasbourg, Illkirch, France) and dissolved in 100% DMSO to a concentration of 50 mM. An intermediate stock of each compound was prepared at 20 mM in 50% DMSO. Dose-response experiments were performed in TCM, starting with a compound concentration of 100 µM and 0.25% DMSO and carrying out eleven two-fold dilutions in 0.25% DMSO. A control in TCM without compound and 0.25% DMSO was always added to the assay. The plates were incubated at 37 °C in the presence of 5% CO_2_ as described above. Cell size was determined after staining with lactofuchsin and analysis with the Cytell microscope was performed following the automatic pipeline described previously.

### Endogenous ROS detection by flow cytometry

To detect endogenous reactive oxygen species (ROS), we use the ROS susceptible probe dihydrofluorescein diacetate (DHF, Sigma-Aldrich), prepared at a stock concentration of 4 mM in 100% DMSO. When this probe reacts with ROS, it produces fluorescein molecules, providing green fluorescence. Yeasts were cultured overnight in Sabouraud liquid medium as described previously, washed with PBS and suspended at 5 x 10^4^ cells/mL in Sabouraud liquid medium or TCM. Cells were incubated in 12-well tissue culture plates (Falcon) at 37 °C with a 5% CO_2_ enriched atmosphere for different times (0, 3, 6 and 24 h). At each time, cells were stained with 40 µM DHF (1/100 dilution from initial stock) for the detection of endogenous reactive oxygen species. The cells were incubated for 30 minutes at 37 °C with 5% CO_2_ and then washed twice with PBS. For each time, a control sample without DHF was carried out. Fluorescence intensity measurement was done in a BD Accuri C6 Plus cytometer (BD Biosciences), counting 10000 events. The data was analysed with FlowJo v10 software (Tree Star, Inc).

As a control for DHF activity, cells were also treated at time 0 h with 1 µg/mL of Amphotericin B for 1 h at 37 °C with 5% CO_2_. Then DHF was added and fluorescence was detected as described above (data not shown).

In some experiments, retinoic acid (25 µM) was added during the incubation of TCM and ROS were detected as described above.

### Study of mitochondrial membrane potential by JC-1

The MitoProbe JC-1 Assay Kit (5’,6,6’-tetrachloro-1,1’,3,3’- tetraethylbenzimidazolylcarbocyanine iodid (JC-1), Life Technologies) was used for the study of mitochondrial membrane potential. A stock at 200 µM was prepared in DMSO. This dye is accumulated in the mitochondria and produces green and red fluorescence depending on the membrane potential. When the mitochondria are functional, JC-1 emits both red and green fluorescence. In contrast, when the mitochondria are depolarized, this probe only emits green fluorescence. In this way, changes in red/green fluorescence ratio are indicators of variations in the mitochondrial membrane potential.

The cells were incubated at a cell density of 5 x 10^4^ cells/mL in Sabouraud liquid medium and TCM at 37 °C with a 5% CO_2_ enriched atmosphere as described in the previous sections. After different incubation times (0, 3, 6 and 24 h), cultures were washed twice with PBS, suspended at 5 x 10^4^ cells/mL and JC-1 was added at a final concentration of 2 µM. The samples were incubated at 37 °C and 5% CO_2_ and fluorescence intensity of the cells was measured by flow cytometry using a BD Accuri C6 Plus cytometer (FL-1 channel for green fluorescence and FL-2 channel for red fluorescence) was read in. Data was analysed with FlowJo v10 software (Tree Star, Inc). To determine the red/green fluorescence ratio, we exported the raw fluorescence data in FlowJo to a .csv document which was processed with Excel. After setting a threshold of 50 to exclude not labelled events, the ratio FL2 (red) / FL1 (green) for each cell was calculated. Then, the data was analysed with GraphPad Prism 9.0 and the geometric mean for each population was calculated.

### Mitotracker staining

Cells were cultivated overnight and then incubated in Sabouraud liquid medium and TCM at 37 °C and 5% CO_2_ as described in the previous sections. The cells were incubated for different times (0, 3, 6 and 24 h), and at each time point, a sample of the culture was collected, washed with PBS and suspended at 5 x 10^4^ cell/mL. MitoTracker Red CMXRos (Invitrogen) was prepared at 1 mM in 100% DMSO and then an intermediate stock of 40 µM in PBS was freshly prepared Mitotracker was added at 40 nM to these samples and incubated for 30 minutes at 37 °C with a 5% CO_2_ enriched atmosphere. Samples were washed twice with PBS and the red fluorescence of the cells was imaged in a Stellaris confocal microscope (Leica Microsystems). Images were processed with Fiji and Adobe Photoshop CS3.

### Transmission Electron Microscopy and quantification of the mitochondria

*C. neoformans* H99 cells were grown overnight as previously described. Cells were cultured to a final density of 5 x 10^4^ cells/mL in Sabouraud liquid medium for regular cells and in TCM for titan-like cells and incubated for 16 h at 37 °C with a 5% CO_2_ enriched atmosphere. Both samples were pelleted down at 3,250 g. The pellet (approximately 30 µL pellet) was washed twice with PBS and gently suspended in 2.5% paraformaldehyde + 0.1% glutaraldehyde in 0.1 M phosphate buffer, pH 7.2. After fixation for 2 h at RT, the cells were washed with PBS and remaining free aldehydes were quenched with 50 mM NH_4_Cl in PBS. The cell pellets were embedded in 12% gelatine in PBS and solidified on ice. Next, cubes of 1 mm^3^ were cut and infiltrated with 2.3 M sucrose in PBS overnight at 4 °C. The cubes were mounted on metal pins and frozen by plunging into liquid nitrogen. Thin sections were cut with a cryo-microtome UC7 (Leica Microsystems) with a nominal thickness of 70 nm and picked up with a 1:2 mixture of 2% methylcellulose in water and 2.3 M sucrose in PBS. After thawing, the sections were deposited on 100 mesh copper grids with a carbon, formvar film. To contrast the grids, they were incubated on PBS for 30 min at 37 °C, washed extensively with water, and incubated for 5 min on ice with 0.4% uranyl acetate in 1.8% methylcellulose in water. Excess of contrasting solution was removed by blotting the grids with filter paper. Images were taken in a Tecnai G2 microscope operated at 120 kV with a Ceta camera.

To measure the surface area of the mitochondria on thin sections, images of all mitochondria from 10 cell profiles were taken at a nominal magnification of 30,000x. The surface area of mitochondria was measured with Fiji software using the freehand selection tool. The number of cristae were counted to calculate the number of cristae per surface area for each mitochondrion. In total 58 mitochondria were analysed for the control cells and 106 for titan-like cells.

### Statistical Analysis

Normality of the samples was assessed by the Kolgomorov-Smirnov test. When the samples presented a Gaussian distribution, statistical differences of the cell sizes were assessed using ANOVA with the Bonferroni post-test (significance when p<0.05). If the samples were not normally distributed, we used the Kruskal-Wallis test with the Dunńst post-test. All calculations were performed using GraphPad Prism 9.0.

## Results

### Determination of the optimal DMSO concentration at which DMSO did not inhibit the formation of titan-like cells

The compounds of the Prestwick Chemical Library are prepared at a concentration of 10 mM in 100% DMSO, so we first evaluated the effect of this solvent on titan-like cell formation. For this purpose, we performed a dose-response curve to determine the maximum DMSO concentration that did not interfere with this process. As shown in Figure 1, *C. neoformans* significantly enlarged in size at all the DMSO concentrations tested. However, we found that at concentrations between 0.5 - 2%, there was a partial inhibition of cell growth. In consequence, we decided that the final concentration of DMSO should not exceed 0.25% during titan-like cell induction.

**Figure 1.**
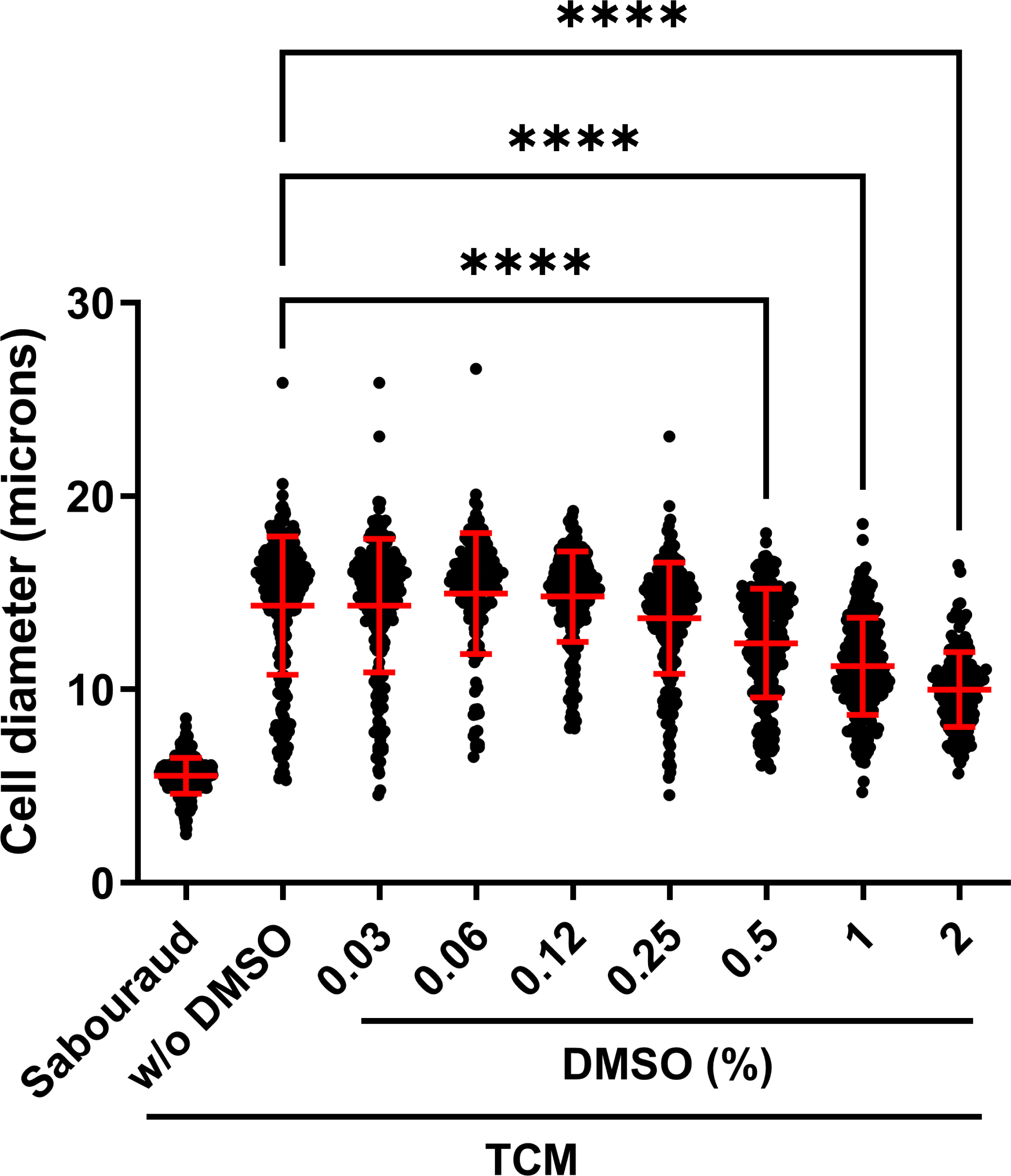
Effect of DMSO in the formation of titan-like cells. A dose- response curve in TCM with different concentrations of DMSO was performed to determine the maximum percentage of DMSO to be used during the screening of the Prestwick Chemical Library. Cells in Sabouraud liquid medium were added as control of regular size cells. Asterisks denote statistical differences (p<0.0001, Kruskal-Wallis test) and the red bars denote the average and the standard deviation of the population.

### Standardization of a fluorescence-based protocol to visualize titan-like cells

We next wanted to describe a protocol that allowed us to visualize the size of the cells based on fluorescence, since this easies further automatic analysis of the images compared to bright field pictures. Our laboratory has extensively employed other dyes to better visualize fungal structures, being lactofuchsin one of the most used. Lactofuchsin binds unspecifically to the cell wall of fungi, allowing its visualization with a light red-pink colour. We observed that it also binds to *C. neoformans*, providing a strong fluorescence using standard rhodamine fluorescence cubes (Figure 2). For this reason, we investigated if lactofuchsin was suitable to visualize the whole size of the cells, including the capsule.

**Figure 2.**
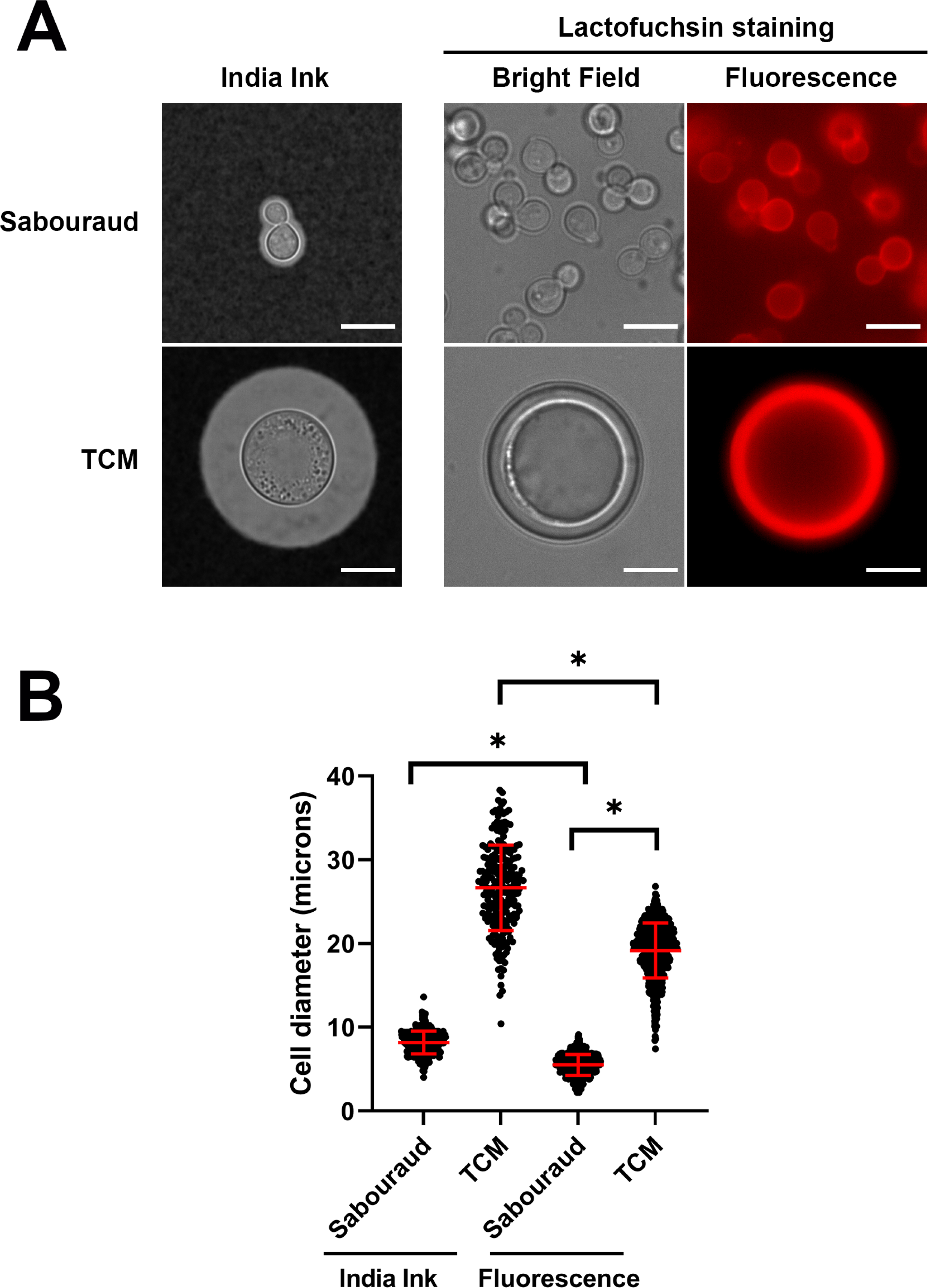
Sizes of C*. neoformans* measured after suspension in India ink or with lactofuchsin staining. A) Morphology of the cells grown in Sabouraud liquid medium (upper panels) and TCM (lower panels) with different stainings, India ink (right) and lactofuchsin (left). B) Differences in size measured after suspension in India ink or staining with lactofuchsin. Asterisks show statistical difference (p<0.0001, t-test) and the red bars denote the average and the standard deviation of the population.

At concentrations around 5 µg/mL, we observed a clear and bright fluorescence staining of titan-like and regular cells, including the capsule. In the case of titan- like cells, this staining was also visible in bright field because the capsule of the cells was visible due to a quellung-effect like phenomenon (see Figure 2). Then, we confirmed that the staining with lactofuchsin did not alter the regular cell size measured by other standard protocols. As shown in Figure 2, the size of the cells detected by lactofuchsin fluorescence was slightly smaller to that measured after suspending the cells in India ink. This was mainly due to an increase in capsular packing, and was even observed when the cells were suspended in lactic acid, the solvent in which fuchsin is dissolved (data not shown). Despite this difference, lactofuchsin staining still clearly differentiated titan-like cells from control cells incubated in non-inducing medium (Figure 2).

### Screening for compounds that inhibit the formation of titan-like cells in *Cryptococcus neoformans*

To identify compounds that inhibited titan-like cell formation, we used the Prestwick Chemical Library, which contains 1,520 off-patent drugs and has been used in repurposing experiments. The compounds in this library are dissolved at 10 mM in 100% DMSO. To avoid the small inhibitory effect of DMSO on titan cell formation at concentrations above 0.5%, for the screening we chose a concentration of 25 µM (1/400 dilution of the original chemical library) for each drug (0.25% DMSO concentration).

For the screening, plates with TCM medium and individual compounds were prepared as described in M&M and inoculated at 10^4^ cells/mL. After an o.n. incubation, the plates were visualized under the microscope to identify wells in which the cells had not grown in size. In addition, we stained them with lactofuchsin and cell size was measured with a Cytell microscope and an in- house developed pipeline that identified and measured all the cell sizes from all the images (see M&M).

We identified 99 compounds that inhibited titan-like cells development. We categorized these compounds depending on the final size of the cells after the incubation in TCM (≤10 microns and >10 - ≤15 microns). Among these, we found several well-known antifungals, such as amphotericin B, voriconazole, itraconazole, fluconazole, nystatine and terbinafine. We also found drugs that presented inhibitory activity against *C. neoformans* in previous screenings performed with the same library (51, 52). For further analysis of the results, we discarded these compounds, finally obtaining 64 potential drugs that inhibited titan-like cell formation. The identified compounds belonged to several therapeutic classes, including antimicrobials, endocrinology, metabolism, central nervous systems, dermatology, allergology, oncology and rheumatology (see table 1 for the complete list).

**Table 1.**
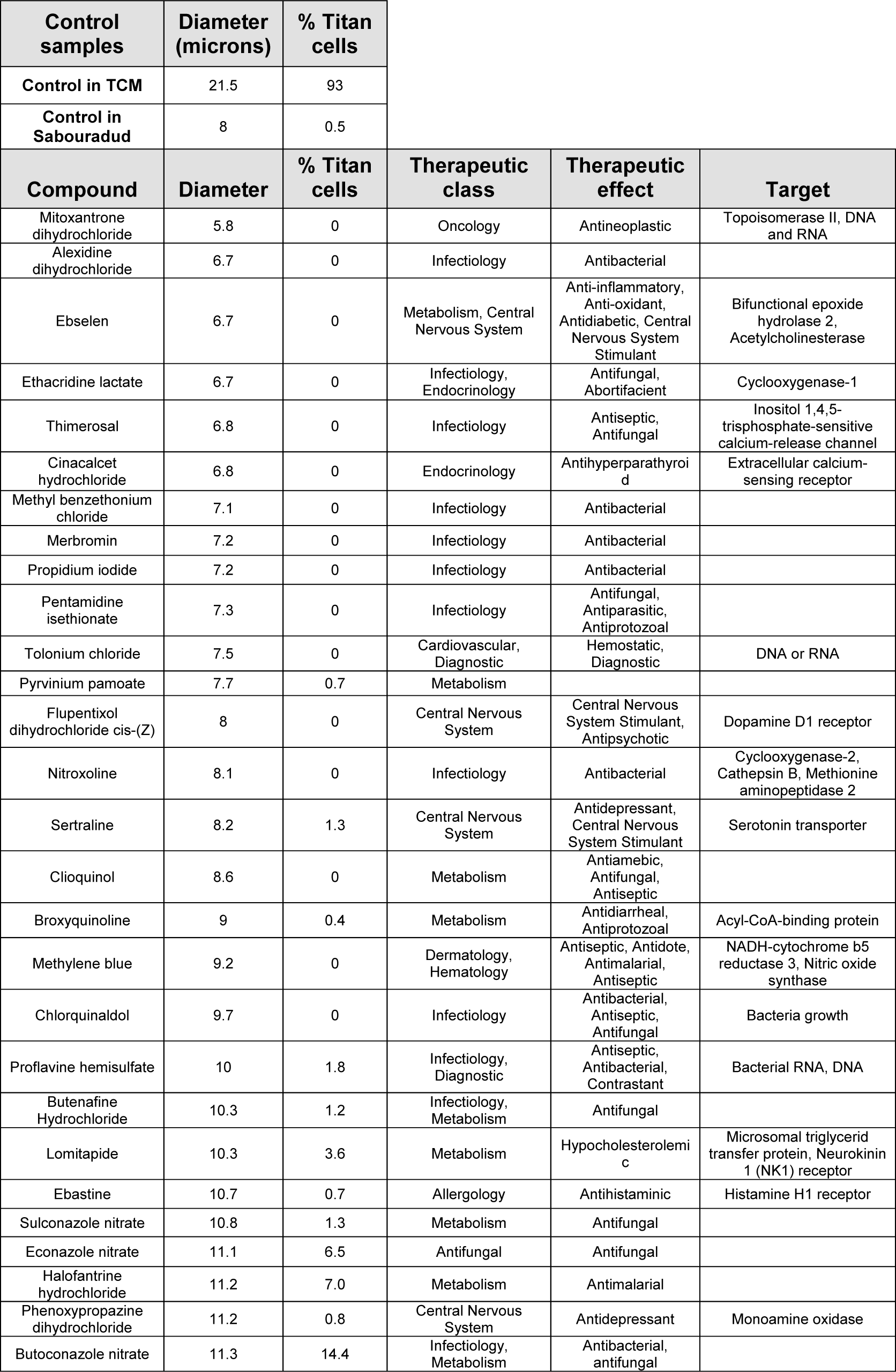
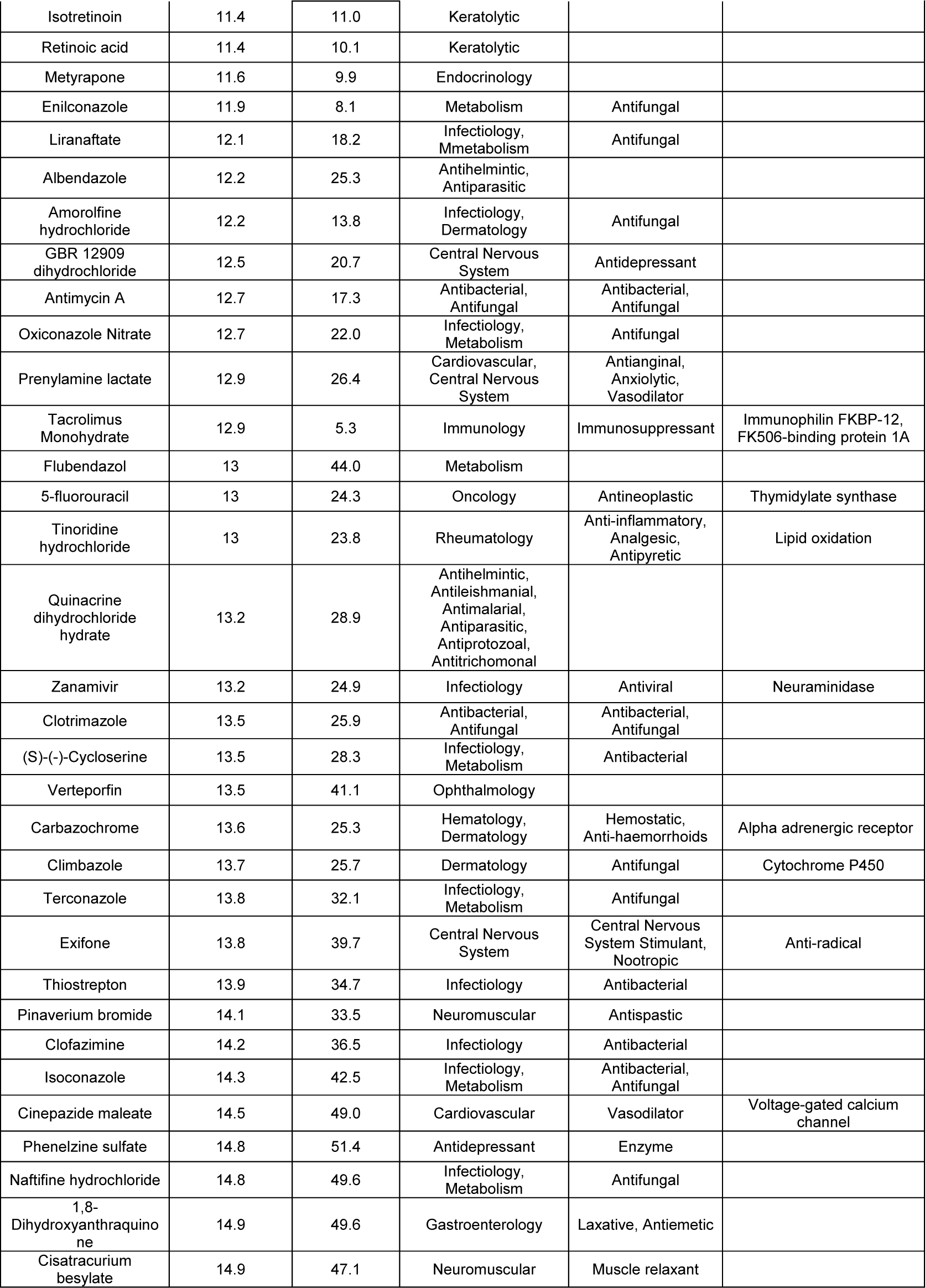
List of compounds that inhibited TC formation.

| Control samples | Diameter (microns) | % Titan cells |  |  |  |
| --- | --- | --- | --- | --- | --- |
| Control in TCM | 21.5 | 93 |  |  |  |
| Control in Sabouradud | 8 | 0.5 |  |  |  |
| Compound | Diameter | % Titan cells | Therapeutic class | Therapeutic effect | Target |
| Mitoxantrone dihydrochloride | 5.8 | 0 | Oncology | Antineoplastic | Topoisomerase II, DNA and RNA |
| Alexidine dihydrochloride | 6.7 | 0 | Infectiology | Antibacterial |  |
| Ebselen | 6.7 | 0 | Metabolism, Central Nervous System | Anti-inflammatory, Anti-oxidant, Antidiabetic, Central Nervous System Stimulant | Bifunctional epoxide hydrolase 2, Acetylcholinesterase |
| Ethacridine lactate | 6.7 | 0 | Infectiology, Endocrinology | Antifungal, Abortifacient | Cyclooxygenase-1 |
| Thimerosal | 6.8 | 0 | Infectiology | Antiseptic, Antifungal | Inositol 1,4,5-trisphosphate-sensitive calcium-release channel |
| Cinacalcet hydrochloride | 6.8 | 0 | Endocrinology | Antihyperparathyroid | Extracellular calcium-sensing receptor |
| Methyl benzethonium chloride | 7.1 | 0 | Infectiology | Antibacterial |  |
| Merbromin | 7.2 | 0 | Infectiology | Antibacterial |  |
| Propidium iodide | 7.2 | 0 | Infectiology | Antibacterial |  |
| Pentamidine isethionate | 7.3 | 0 | Infectiology | Antifungal, Antiparasitic, Antiprotozoal |  |
| Tolonium chloride | 7.5 | 0 | Cardiovascular, Diagnostic | Hemostatic, Diagnostic | DNA or RNA |
| Pyrrinium pamoate | 7.7 | 0.7 | Metabolism |  |  |
| Flupentixol dihydrochloride cis-(Z) | 8 | 0 | Central Nervous System | Central Nervous System Stimulant, Antipsychotic | Dopamine D1 receptor |
| Nitroxoline | 8.1 | 0 | Infectiology | Antibacterial | Cyclooxygenase-2, Cathepsin B, Methionine aminopeptidase 2 |
| Sertraline | 8.2 | 1.3 | Central Nervous System | Antidepressant, Central Nervous System Stimulant | Serotonin transporter |
| Clioquinol | 8.6 | 0 | Metabolism | Antiamoebic, Antifungal, Antiseptic |  |
| Broxyquinoline | 9 | 0.4 | Metabolism | Antidiarrheal, Antiprotozoal | Acyl-CoA-binding protein |
| Methylene blue | 9.2 | 0 | Dermatology, Hematology | Antiseptic, Antidote, Antimalarial, Antiseptic | NADH-cytochrome b5 reductase 3, Nitric oxide synthase |
| Chlorquinaldol | 9.7 | 0 | Infectiology | Antibacterial, Antiseptic, Antifungal | Bacteria growth |
| Proflavine hemisulfate | 10 | 1.8 | Infectiology, Diagnostic | Antiseptic, Antibacterial, Contrastant | Bacterial RNA, DNA |
| Butenafine Hydrochloride | 10.3 | 1.2 | Infectiology, Metabolism | Antifungal |  |
| Lomitapide | 10.3 | 3.6 | Metabolism | Hypocholesterolemic | Microsomal triglycerid transfer protein, Neurokinin 1 (NK1) receptor |
| Ebastine | 10.7 | 0.7 | Allergology | Antihistaminic | Histamine H1 receptor |
| Sulconazole nitrate | 10.8 | 1.3 | Metabolism | Antifungal |  |
| Econazole nitrate | 11.1 | 6.5 | Antifungal | Antifungal |  |
| Halofantrine hydrochloride | 11.2 | 7.0 | Metabolism | Antimalarial |  |
| Phenoxypropazine dihydrochloride | 11.2 | 0.8 | Central Nervous System | Antidepressant | Monoamine oxidase |
| Butoconazole nitrate | 11.3 | 14.4 | Infectiology, Metabolism | Antibacterial, antifungal |  |
| Isotretinoin | 11.4 | 11.0 | Keratolytic |  |  |
| Retinoic acid | 11.4 | 10.1 | Keratolytic |  |  |
| Metirapone | 11.6 | 9.9 | Endocrinology |  |  |
| Enilconazole | 11.9 | 8.1 | Metabolism | Antifungal |  |
| Liranaftate | 12.1 | 18.2 | Infectiology,<br>Mmetabolism | Antifungal |  |
| Albendazole | 12.2 | 25.3 | Anthelmintic,<br>Antiparasitic |  |  |
| Amorolfine<br>hydrochloride | 12.2 | 13.8 | Infectiology,<br>Dermatology | Antifungal |  |
| GBR 12909<br>dihydrochloride | 12.5 | 20.7 | Central Nervous<br>System | Antidepressant |  |
| Antimycin A | 12.7 | 17.3 | Antibacterial,<br>Antifungal | Antibacterial,<br>Antifungal |  |
| Oxiconazole Nitrate | 12.7 | 22.0 | Infectiology,<br>Metabolism | Antifungal |  |
| Prenylamine lactate | 12.9 | 26.4 | Cardiovascular,<br>Central Nervous<br>System | Antianginal,<br>Anxiolytic,<br>Vasodilator |  |
| Tacrolimus<br>Monohydrate | 12.9 | 5.3 | Immunology | Immunosuppressant | Immunophilin FKBP-12,<br>FK506-binding protein 1A |
| Flubendazol | 13 | 44.0 | Metabolism |  |  |
| 5-fluorouracil | 13 | 24.3 | Oncology | Antineoplastic | Thymidylate synthase |
| Tinoridine<br>hydrochloride | 13 | 23.8 | Rheumatology | Anti-inflammatory,<br>Analgesic,<br>Antipyretic | Lipid oxidation |
| Quinacrine<br>dihydrochloride<br>hydrate | 13.2 | 28.9 | Anthelmintic,<br>Antileishmanial,<br>Antimalarial,<br>Antiparasitic,<br>Antiprotozoal,<br>Antitrichomonal |  |  |
| Zanamivir | 13.2 | 24.9 | Infectiology | Antiviral | Neuraminidase |
| Clotrimazole | 13.5 | 25.9 | Antibacterial,<br>Antifungal | Antibacterial,<br>Antifungal |  |
| (S)-(-)-Cycloserine | 13.5 | 28.3 | Infectiology,<br>Metabolism | Antibacterial |  |
| Verteporfin | 13.5 | 41.1 | Ophthalmology |  |  |
| Carbazochrome | 13.6 | 25.3 | Hematology,<br>Dermatology | Hemostatic,<br>Anti-haemorrhoids | Alpha adrenergic receptor |
| Climbazole | 13.7 | 25.7 | Dermatology | Antifungal | Cytochrome P450 |
| Terconazole | 13.8 | 32.1 | Infectiology,<br>Metabolism | Antifungal |  |
| Exifone | 13.8 | 39.7 | Central Nervous<br>System | Central Nervous<br>System Stimulant,<br>Nootropic | Anti-radical |
| Thiostrepton | 13.9 | 34.7 | Infectiology | Antibacterial |  |
| Pinaverium bromide | 14.1 | 33.5 | Neuromuscular | Antispastic |  |
| Clofazimine | 14.2 | 36.5 | Infectiology | Antibacterial |  |
| Isoconazole | 14.3 | 42.5 | Infectiology,<br>Metabolism | Antibacterial,<br>Antifungal |  |
| Cinepazide maleate | 14.5 | 49.0 | Cardiovascular | Vasodilator | Voltage-gated calcium<br>channel |
| Phenelzine sulfate | 14.8 | 51.4 | Antidepressant | Enzyme |  |
| Naftifine hydrochloride | 14.8 | 49.6 | Infectiology,<br>Metabolism | Antifungal |  |
| 1,8-Dihydroxyanthraquino<br>ne | 14.9 | 49.6 | Gastroenterology | Laxative, Antiemetic |  |
| Cisatracurium<br>besylate | 14.9 | 47.1 | Neuromuscular | Muscle relaxant |  |

### Dose-response curve of selected compounds

To validate the results of the assay, we selected 10 compounds based on their mechanism of action and effect and performed dose-response experiments (from 100 to 0.1 µM, with a constant concentration of 0.25% of DMSO). These compounds were isotretinoin, retinoic acid, mitoxantrone dihydrochloride, pentamidine isethionate, alexidine dihydrochloride, clioquinol, metyrapone, sertraline, ebselen and antimycin A. As shown in Figure 3, we confirmed that all selected compounds inhibited titan cell formation in a dose-dependent manner. Most of the drugs showed a lineal inhibition between 6-100 µM. In the case of mitoxantrone, concentrations around 6 µM fully inhibited cryptococcal cell growth. Finally, alexidine dihydrochloride was the compound that showed the strongest inhibitory effect with concentrations below 1 µM, blocking almost completely the formation of titan cells.

**Figure 3.**
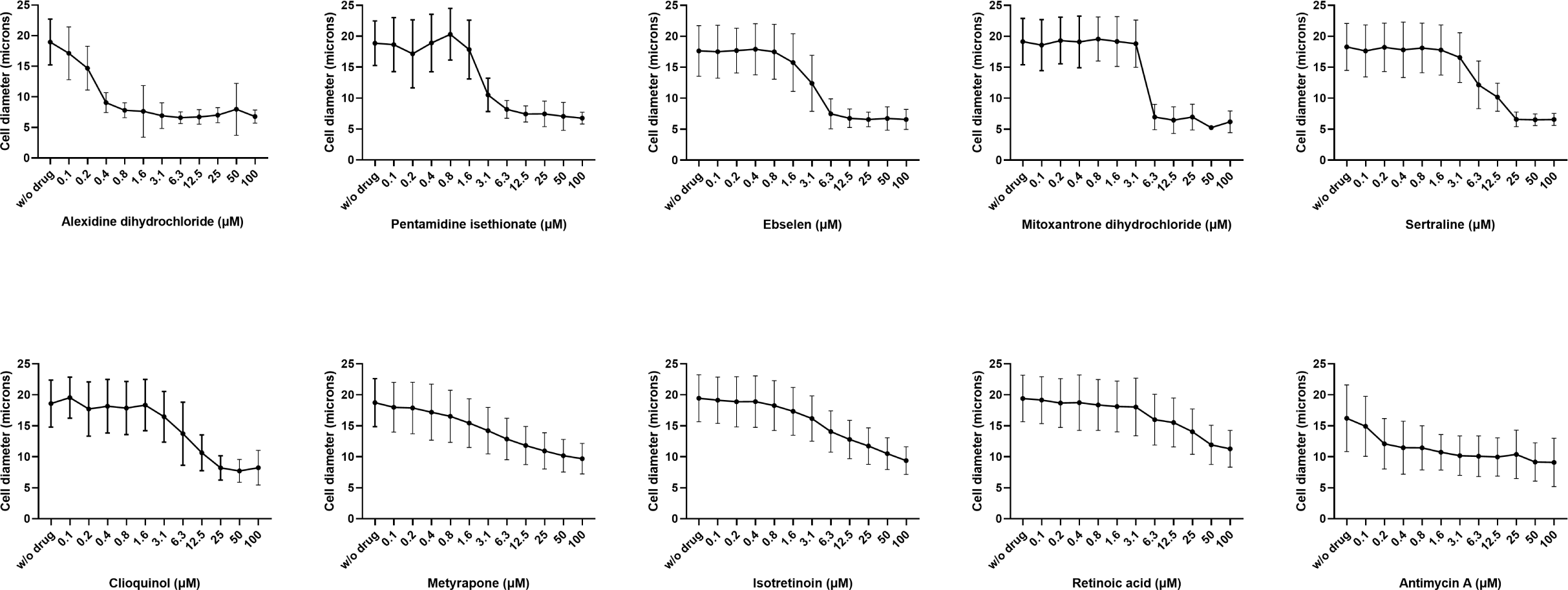
Dose-response curves of the selected compounds. Cells were incubated in TCM in the presence of different concentrations of each compound (ranging from 100 µM to 0.1 µM) as described in M&M. A control without compound was added in all curves. Cells were stained with lactofuchsin and imaged with a Cytell microscope (see M&M). Diameters of the cells were measured automatically and the average of the cell sizes and the standard deviations are represented in the figure. The experiment was performed in duplicate.

### Detection of reactive oxygen species and mitochondrial membrane potential during titan-like cell formation

We observed that some compounds from our list are well characterized antioxidants, such as retinoic acid, isotretinoin (or 13-cis retinoic acid) and ebselen. Interestingly, retinoic acid and isotretinoin had almost the same inhibitory profile. In consequence, we hypothesized that during titanization, an endogenous accumulation of free radicals could trigger a stress signal required for titan-like cell formation. To test this hypothesis, we examined if during this process there was an accumulation of endogenous reactive oxygen species (ROS). We used a ROS-sensitive probe (dihydrofluorescein, DHF), which is fluorescent in presence of free radicals. With this probe, we estimated the production of free radicals in titan cell inducing medium (TCM) or in media where the cells do not increase in size (Sabouraud). As shown in Figure 4, we found that during incubation in TCM, there was a gradual accumulation of ROS in the cells, already noticeable at 3 and 6 h, and all the cells showed a strong fluorescence signal at 24 h. In contrast, the increase of intracellular ROS in non- inducing conditions was not significant at short times (3 and 6 h), and only after 24 h of incubation the cells presented a significant fluorescence signal, although it was lower than the peak observed in TCM. This result indicated that during titan-like cell formation there is a significant intracellular accumulation of ROS.

**Figure 4.**
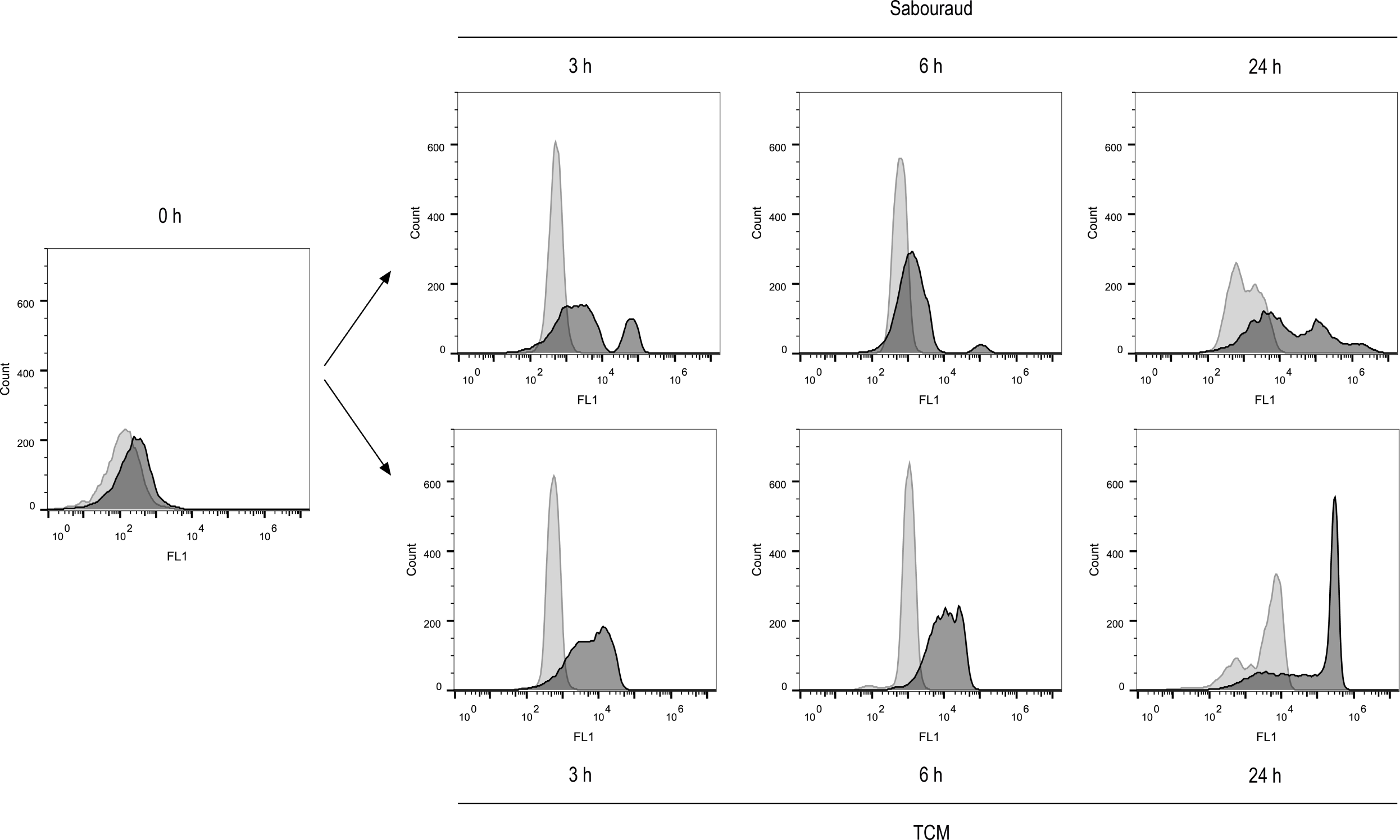
Endogenous ROS detection by dihydrofluorescein diacetate (DHF) during titan-like cells induction. Cells from H99 strain were incubated o.n. in Sabouraud liquid medium and then transferred to the same medium (as control of regular cells) or to titan cell medium (TCM). ROS were measured at 0, 3, 6 and 24 h after addition of DHF (dark grey histogram) and detection by flow cytometry. A parallel sample without DHF (light grey histogram) was carried out in parallel. A control with AmB 1µg/mL was added at time 0 h to confirm ROS detection by DHF (data not shown). The experiments were performed twice in two different days.

We next investigated if the addition of retinoic acid had any effect on the accumulation of ROS in cells incubated in TCM. As shown in Figure 5, addition of the antioxidant reduced the amount of endogenous free radicals accumulated during titanization.

**Figure 5.**
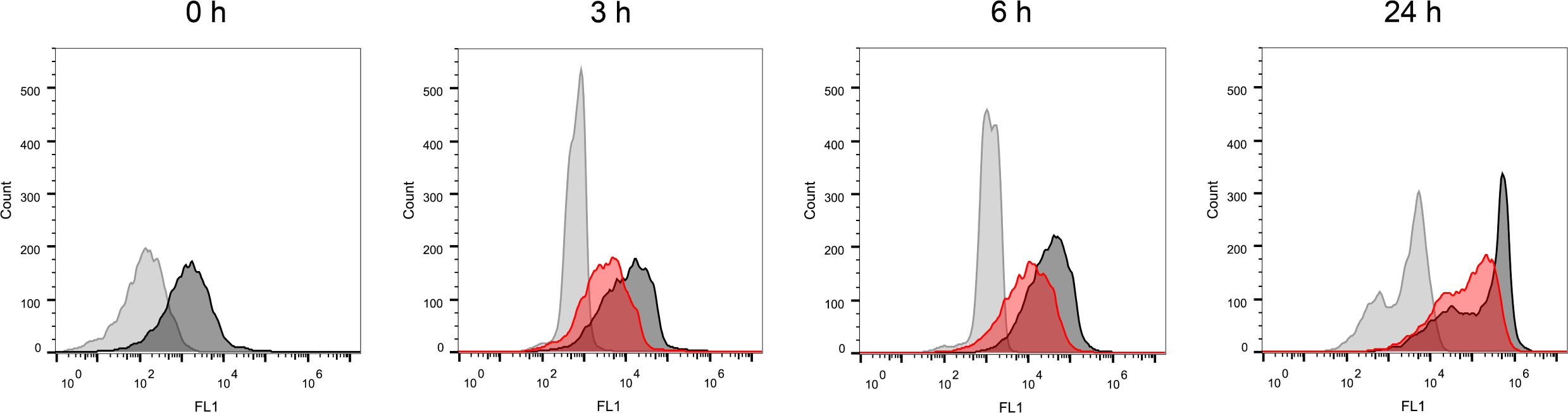
Effect of retinoic acid on ROS production during titanization. Titan-like cells were obtained and stained with DHF to measure accumulation of ROS as described in Figure 4. Retinoic acid at 25 µM (light red histogram) was added at time 0 h to all samples and ROS production was detected by the addition of 40 µm DHF at time 0, 3, 6 and 24 h (dark grey histogram). A control without retinoic acid and without DHF (light grey histogram) was also measured at each time points.

Since free radicals are mainly produced in mitochondria as subproducts of respiratory electron transport, we evaluated functionality of this organelle with several complementary approaches. We first measured if there was any variation in the mitochondrial membrane potential using the fluorescent probe JC-1 (see M&M). This probe can emit both red and green fluorescence, and membrane depolarization is characterized by a decrease in the red/green fluorescence ratio. We added this probe after 0, 3, 6 and 24 hours of incubation in TCM and Sabouraud and measured both the red and green fluorescence signal of the cells. We found that the staining was different in both mediums. In TCM there was a clear increase of both red and green fluorescence signals of the cells after 3 and 6 h of incubation (Figure 6). This increase was not noticeable in the non-inducing conditions. The stronger staining suggested that there was an increase in mitochondrial mass during titanization. In addition, we noticed that during titan-like cells formation there were different populations with different red/green fluorescence ratios, and analysed them using different gates. We found that there was a partial depolarization of the mitochondria in both conditions (see table 2). Control cells incubated in Sabouraud medium had a similar ratio than the cells in TCM from gate 1 after 3 and 6 h of incubation. In contrast, the smaller population of cells from gate 2 had a lower ratio. After 24 h of incubation, the ratio of the fluorescence was higher in titan-like cells than in those cultivated in Sabouraud medium, indicating that in these titan-like cells there is not only more mitochondrial mass, but also a higher activity than in cells of regular size.

**Figure 6.**
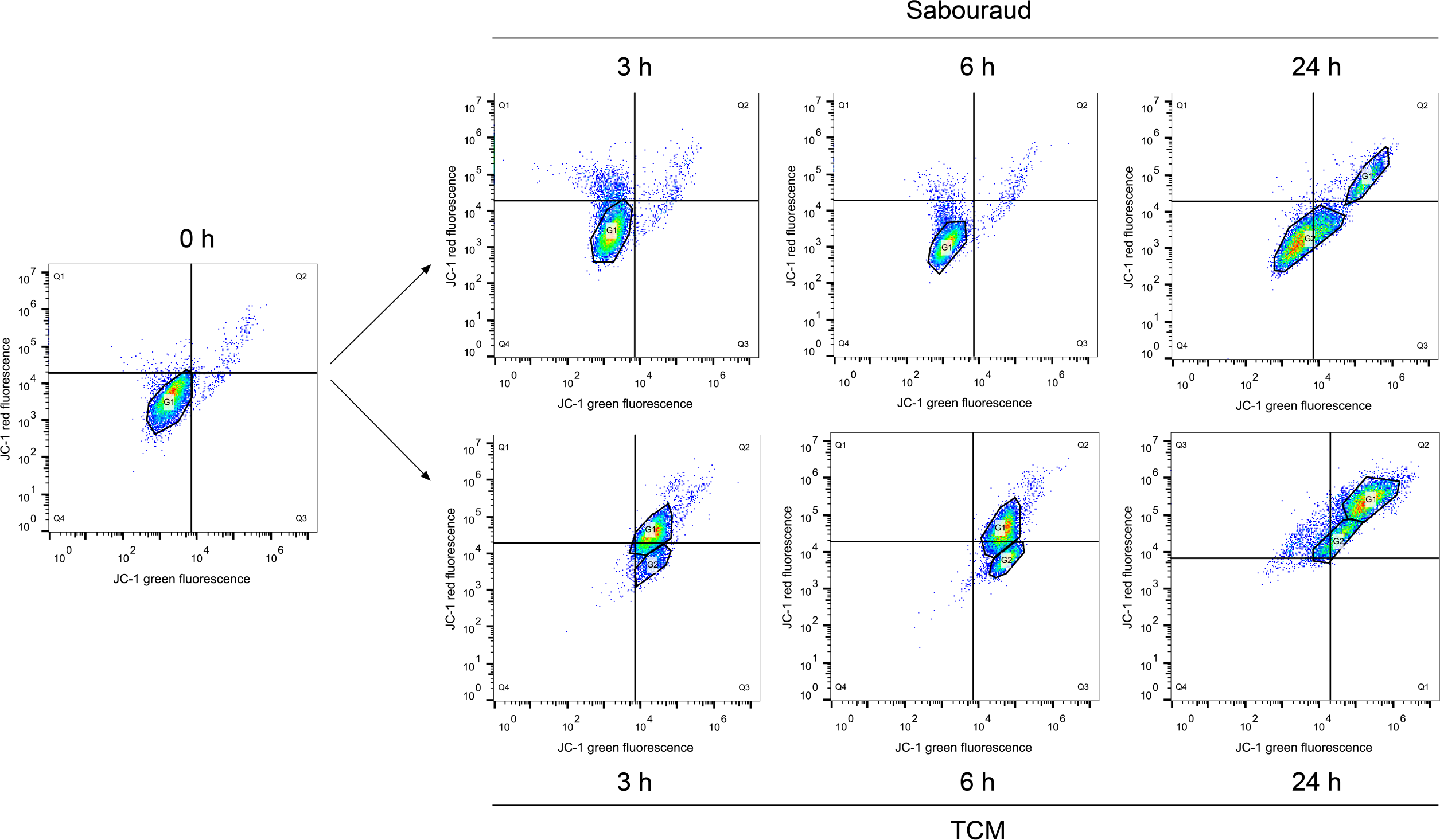
Determination of membrane potential by JC-1 during titan cell- like induction. *C. neoformans* H99 cells were grown o.n. in Sabouraud liquid medium and then transferred to the same medium (as control of regular cells) and to titan cell medium (TCM) (see M&M). Mitochondrial membrane potential was measured at 0, 3, 6 and 24 h by flow cytometry after addition of JC-1 and measurement of green (FL-1 channel, x-axis) and red (FL-2 channel, *y*-axis) fluorescence signals was carried out. The sample of Sabouraud cells at time 0 h was used to determine the quadrants. At some time points, specific gates (denoted by a polygon) were defined for each dot plot to analyse specific cell populations. The experiment was performed in duplicates in two different days.

**Table 2.** Red / Green fluorescence intensity ratios after JC-1 staining during titan-like cell formation. The fluorescence intensity from cells from the cytometry experiment from Figure 5 was analysed in Excel and Graphpad, and the ratio of the Red/Green fluorescence intensity was obtained. We calculated the geometric mean of the whole population (numbers in bold) and of the specific gates (G1 and G2) defined in the graph (numbers in italics).

|  | JC-1 Red/Green fluorescence ratio |  |
| --- | --- | --- |
|  | Sabouraud | TCM |
| 0 h | **1.96** | **1.96** |
| 3 h | **1.42** | **1.35** |
|  |  | (G1 – 1.60)<br>(G2 – 0.42) |
| 6 h | **1** | **0.62** |
|  |  | (G1 – 0.91)<br>(G2 – 0.25) |
| 24 h | **0.58** | **0.92** |
|  | (G1 – 0.50)<br>(G2 – 0.58) | (G1 – 1.48)<br>(G2 – 0.97) |

**Table 3.** Differences in the mitochondria of cells cultured in Sabouraud medium and titan cell medium (TCM). The size and the number of cristae of the mitochondria identified by electron microscopy was quantified (n > 40 for each condition), and the average ± standard deviation of each parameter is shown in the table. Statistical differences were assessed using the Kruskal-Wallis test with Dunńs post-test for multiple comparisons.

|  | Sabouraud | TCM | Ratio<br>(TCM/Sab) | p-value |
| --- | --- | --- | --- | --- |
| Mitochondrial area ($\mu\text{m}^2$) | 0.175 ± 0.131 | 0.213 ± 0.114 | 1.22 | < 0.0001 |
| N of cristae / mitochondria | 3.7 ± 1.9 | 9.1 ± 4.5 | 2.5 | < 0.0001 |
| N of cristae / mitochondrial area ($\mu\text{m}^2$) | 25.6 ± 13.9 | 46.3 ± 16.8 | 1.8 | < 0.0001 |

### Mitotracker staining

To visualize mitochondrial network organization during titanization, we used mitotracker dye. In *C. neoformans,* mitochondria can accumulate in a fragmented, tubular or diffuse pattern (63). Cells incubated in Sabouraud medium provided mainly a fragmented pattern after mitotracker staining. In contrast, during titan-like cells development, the mitochondria adopted mainly a clear tubular pattern (Figure 7).

**Figure 7.**
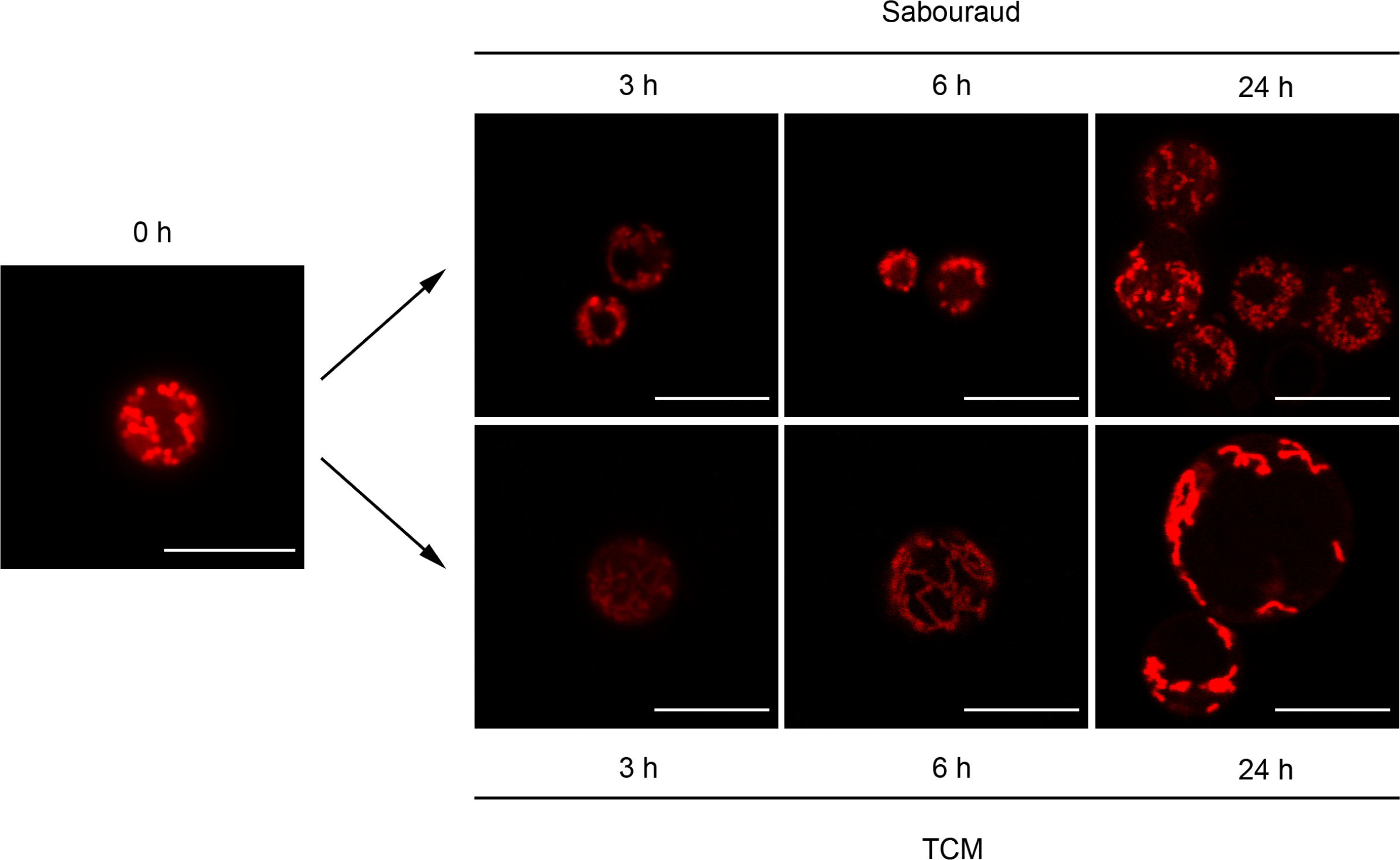
Mitochondrial morphology of *C. neoformans* with MitoTracker Red CMXRos probe. Mitochondrial organization was observed by mitotracker staining (see M&M). Cells were cultivated in Sabouraud or TCM as described above, and mitotracker was added at each time point (0, 3, 6 and 24 h). Fluorescence pattern was observed in a Stellaris confocal microscope (upper row, cells incubated in Sabouraud; lower row, cells incubated in TCM). Scale bar in every panel corresponds to 10 µm.

### Electron microscopy of the mitochondria

Due to the physiological and morphological changes of mitochondria observed by light microscopy, we performed transmission electron microscopy and compared the ultrastructure of this organelle in titan-like cells and regular cells. As shown in Figure 8, the cells presented a large vacuole, which has already been described in these cells (42). We also observed that sample preparation for the analysis of intracellular organelles resulted in partial detachment of cell wall and capsular components (Figure 8A and B). In contrast, the morphology of the mitochondria was well preserved (Figure 8C and D). We found that the mitochondria of titan-like cells presented a significant larger size than those from control cells. Furthermore, there was a striking difference in the number of visible cristae inside the mitochondria. In particular, titan-like cells had the matrix of mitochondria compacted with cristae, which was not observed in control cells. In addition, the cristae of control cells had regular spacing between their membranes, whereas the cristae in the titan like cells showed a swollen, more irregular organization. We quantified the size of the mitochondria and number of cristae. The average size of the mitochondria from titan-like cells was around 1.2 larger than those from control cells, but the number of cristae and cristae/µm^2^ was almost doubled in titan-like cells (Figure 8E, F and G).

**Figure 8.**
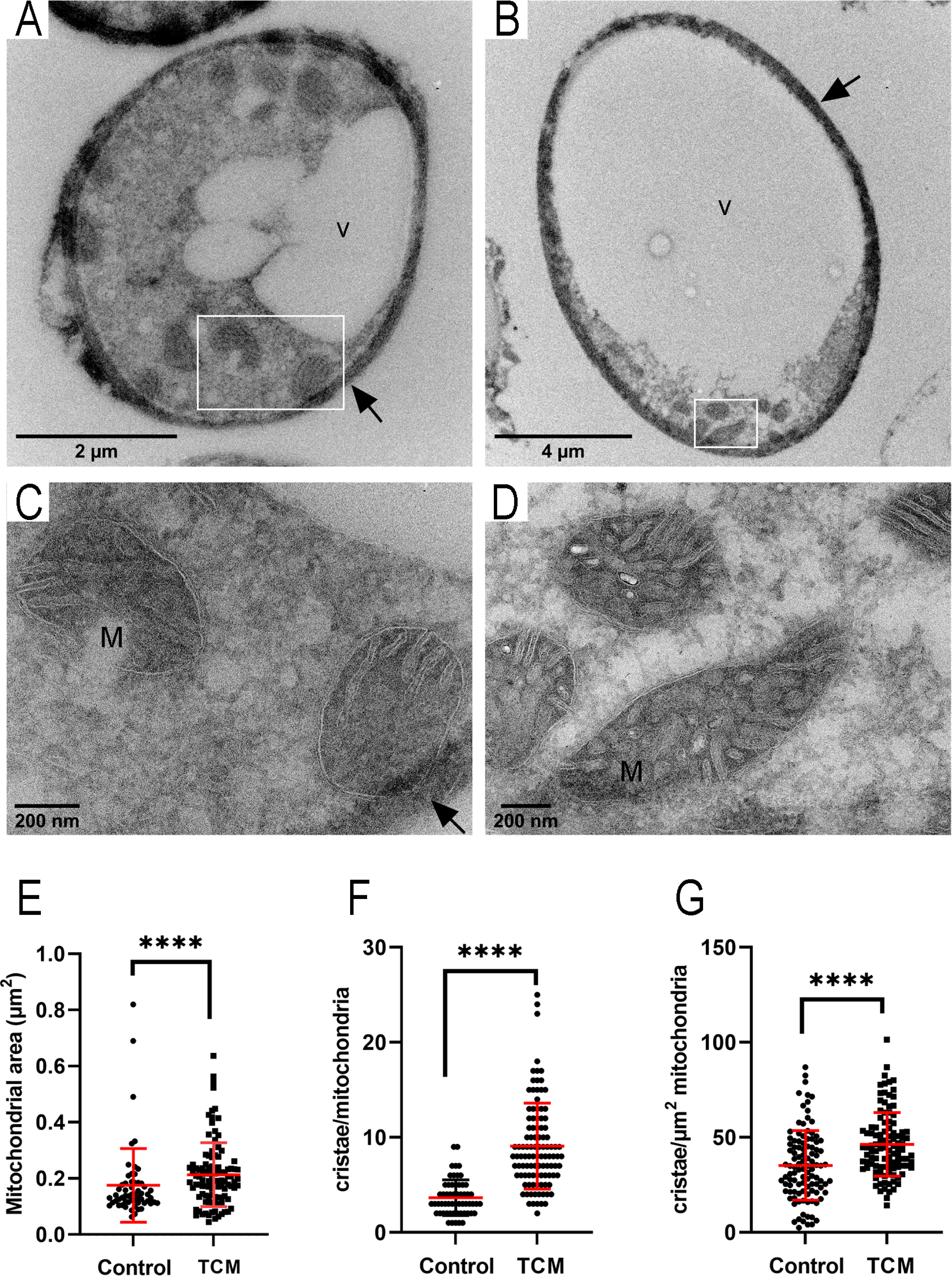
Mitochondrial morphology by transmission electron microscopy. A. Overview image of a regular cell incubated in Sabouraud liquid medium. B. overview image of a titan-like cell cultivated in TCM. White square shows the mitochondria, shown in C and D, respectively. Arrows indicate cell wall detachment. M: mitochondria, V: vacuole. C-D. Higher magnification images of the morphology of mitochondria in regular cells, Sabouraud (C) and titan-like cells (D). The area of each mitochondrium (E), number of cristae per mitochondrium (F) and number of cristae per µm^2^ of mitochondrium (G) were quantified and shown in dot-plot graphs. Asterisks show statistical differences (****, p<0.0001, non-parametric Mann-Whitney t-test).

## Discussion

The possibility to obtain titan-like cells *in vitro* has allowed identifying new inducing factors, signalling pathways and positive and negative regulators of titanization. To obtain new insights about the processes involved in titan cell development, we have used a different approach in which we identified drugs that inhibit this process. For this purpose, we performed a screening using an off-patent library. Since many of the compounds have defined structure and target, we hypothesized that this approach could provide insights on the molecular mechanisms that are required to produce titan cells.

During the standardization of the protocol, we observed that subinhibitory concentrations of DMSO (0.5%-1%) partially blocked the induction of titan cells. This solvent can alter the non-convalent bonds that attach the fibbers of the capsule (64–66), so it is possible that titan cell formation requires an intact capsule structure to attach new polysaccharide fibbers, although we cannot discard that DMSO induces other cell alterations that interferes with this process.

To carry out our screening, we designed a method based on fluorescence as it offers significant advances to perform automatic image analysis. We took advantage of the unspecific binding of the dye lactofuchsin to the surface of fungi. When we compared the binding of this dye with the classical India ink staining, we observed that the size of the cells measured with lactofuchsin was slightly smaller than the one measured with India ink. However, even with this limitation, lactofuchsin staining was still useful to differentiate cells of different size, in particular those incubated in Sabouraud and in the inducing medium TCM. This method offered several advantages, such as fluorescence intensity and low cost. Our protocol allowed an automatic method of image acquisition and bioinformatic analysis that resulted in a detailed analysis of the exact inhibition of each compound.

We identified around 100 inhibitory compounds. Some of them had already been identified in the literature as drugs with antifungal activity against *C. neoformans*. Still, we found more than sixty compounds that selectively inhibited titan cell development. The diversity of compounds we identified reflects that the induction of TC depends on the induction of multiple and overlapping mechanisms and pathways, which is in agreement with previous findings (47–49).

Some of the inhibitory drugs have already defined mechanisms of action. For example, mitoxantrone is an antitumoral that inhibits the activity of topoisomerase II, and affects DNA replication. At the moment we do not know which is the mechanism by which mitoxantrone inhibits TC formation, but the fact that it was not identified in previous screenings suggests that it could exert its effect through a mechanism independent of inhibition of DNA replication. In this sense, this compound inhibits the activity of a mitochondrial calcium importer (67), which has a putative homolog in *C. neoformans* (CNAG_00107), suggesting that mitoxantrone could also interfere through alteration of mitochondrial activity. In agreement, several of the identified compounds also have effects in this organelle. One of them was antimycin A, which is a well- known inhibitor of complex III from the respiratory chain. Other drugs that have other characterized targets also have effects on the mitochondria. For example, pentamidine, that has multiple parasitic effects, can also inhibit mitochondrial topoisomerases and causes a reduction in ATP concentration in *Pneumocystis* and *Tripanosoma* (68–70). Metyrapone, inhibits the steroid 11β-hydroxylase, which is a P450 cytochrome located at the inner membrane of the mitochondria. Furthermore, we found another compound (climbazole) that also inhibits the activity of a cytochrome P450 (71).

Alexidine dihydrochloride was one of the compounds that inhibited TC formation more effectively. This drug blocks bacterial growth by inhibiting the production of extracellular polysaccharide and decreasing adhesion. Capsular components from *C. neoformans* are also released to the medium (72, 73), so it is also possible that alexidine interferes with the trafficking of capsular polysaccharides and in consequence, interferes with titanization. However, this drug has also other action mechanisms, such as inhibition of the mitochondrial phosphatase PTPMT1 (74) and the activity of secreted phospholipases (75).

One striking and unexpected finding of our work was that several compounds that act as antioxidants inhibited the formation of titan cells in *C. neoformans*. This was the case of retinoic acid and its isoform isotriteonin. Ebselen was another compound that inhibited TC formation, and it also has strong antioxidant properties in the cells (76, 77). Based on this result we suggested that ROS are required to induce this process. However, ebselen showed a stronger inhibition compared with the other antioxidants, suggesting that it might exert its inhibitory effect through other mechanisms. We observed with ROS- sensitive fluorescent probes an endogenous accumulation of this type of free radicals. ROS are mainly produced in mitochondria as subproducts of the respiratory chain, so we investigated if there were any functional changes in this organelle. We found that there was a significant increase in JC-1 staining, a change in mitochondrial organization to a tubular, network-like pattern and an increase in size and in the number of intramitochondrial cristae. It has been described that a tubular organization of the mitochondria is achieved after fusion of this organelle and is associated with resistance to stress conditions and virulence (63, 78, 79). Altogether, our data indicates that TC formation is associated to noticeable changes in mitochondrial activity with a subsequent production of ROS, and that this increase in free radicals is required to increase cell size in *C. neoformans*.

ROS concentration in the cell is determined by the balance between its synthesis and detoxification by the antioxidant mechanisms of the cell. Titan cell production requires a significant synthesis of new cellular and capsular components, therefore the cells also need to increase their mitochondrial activity to produce the energy required for this processes. The finding that some compounds that alter the mitochondrial activity interfere with TC formation supports this idea. In addition, we argued that the increase in mitochondrial activity is the most probable explanation that supports the increase in intracellular ROS observed in this work. Noteworthy, *C. neoformans* is a respiratory yeast, so most of the energy production depends on the mitochondrial activity. Furthermore, titanization was induced in this work in a medium that has a low nutrient concentration, so energy supply is most likely obtained by the degradation of internal resources. However, it cannot be discarded that the stress caused by the limitation of nutrients induces a stress signal that leads to ROS accumulation. In this sense, many fungi adapt their morphology between filaments, hyphae and pseudohyphae, and these changes are normally a response to a stress situation, such as high temperature, nutrient starvation or serum. In agreement, titan-like cell formation could represent one of the morphological changes induced in stressful environments.

Our work highlights the importance of endogenous ROS during titan cell formation. ROS play multiple roles in the cell depending on their concentration. A high concentration of free radicals can produce cellular damage on DNA, lipids and proteins, all of them involved in apoptosis. Furthermore, some antifungal compounds, such as amphotericin B, induce a strong oxidative burst in the cells (80–83), participating in the fungicidal effect of this drug. But ROS have also been involved in signalling pathways (84–86), with beneficial effects in some conditions and participating in cellular homeostasis. ROS can also induce authophagy (87), which is used by the cell to obtain energy from fungal compartments. In the case of TC, the possibility that ROS induce autophagy to obtain energy to trigger cell growth is challenging and deserves more detailed studies in the future.

Our findings are also in agreement with a recent report that also highlights the importance of reactive nitrogen species (RNS) and ROS in TC formation (88). These authors also demonstrate that treatment of the cells with a SOD mimetic drug (MnTBAP) can also block titanization, a finding that supports the results presented in our work.

Our results might also provide insights about the mechanism of titan cell formation *in vivo*. During infection, the host elicits several responses and antimicrobial mechanisms, which include the production of ROS by phagocytic cells. It is then possible that this oxidative burst triggers the signalling pathways that induce cryptococcal cell growth and further adaptation to the host environment.

In the context of titan cell appearance in the host, we would also like to highlight that the identification of inhibitory compounds might have therapeutic potential. Titan cells are difficult to eliminate from the lungs and offer a selective advantage to the pathogen to persist during long periods. Inhibition of this process might not facilitate killing of fungal cells by the immune system, but it could also increase the activity of fungal drugs and augment the efficacy of antifungal therapy.

In conclusion, our work has provided a significant number of compounds that inhibit the formation of titan-like cells. The future characterization of the mechanisms of action of these compounds by the scientific community will be a great contribution to understand the molecular mechanisms required for this process, and to design new strategies aim to decrease fungal adaptation in the host. The findings that during TC formation there is a significant increase of mitochondrial activity and accumulation of ROS, together with the inhibitory role of antioxidants in this morphological transition lead us to establish the hypothesis that ROS are key molecules required for TC development in *C. neoformans*. This opens new perspectives on the molecular mechanisms involved in the formation of TC, and will allow new approaches to characterize these cells in the future.

## Acknowledgements

Acknowledgement and funding

This work was funded by projects from the Spanish Ministry for Science and Innovation (SAF2017-86912-R and PID2020-114546RB-I00) to Oscar Zaragoza. Irene García Barbazán was funded by the ministry for Science and Innovation (contract FPI PRE2018-083436). Rocío García-Rodas was funded by a “Juan de la Cierva” Contract from the Spanish Ministry for Economics, Industry and Competitivity (reference: IJCI-2015-25683).

